# Plasmin-mediated cleavage of GPIbα contributes to breakdown of platelet-von Willebrand factor complexes

**DOI:** 10.64898/2026.03.24.713874

**Authors:** Rowan Frunt, Elfie I. Moesker, Kazuya Sakai, Masanori Matsumoto, Albert Huisman, Claudia Tersteeg, Hinde El Otmani

**Affiliations:** Central Diagnostic Laboratory Research, University Medical Center Utrecht, Utrecht University, Utrecht, The Netherlands; Department of Blood Transfusion Medicine, Nara Medical University, Kashihara, Japan; Laboratory for Thrombosis Research, KU Leuven Campus Kulak Kortrijk, Kortrijk, Belgium

## Abstract

Von Willebrand factor (VWF) is an essential contributor to hemostasis through its interaction with the platelet glycoprotein (GP) Ibα receptor. VWF is cleaved by ADAMTS13 to limit its prothrombotic properties. Failure to do so can result in platelet-VWF complexes that occlude the microcirculation, as seen in thrombotic thrombocytopenic purpura (TTP). In this setting, plasmin becomes active to cleave VWF, forming a distinct plasmin-generated cleavage product of VWF (cVWF) that is detectable during acute attacks in patients with TTP and following therapeutic plasminogen activation in a mouse model of TTP. However, it remains unclear whether plasmin-mediated proteolysis of VWF alone accounts for the breakdown of platelet-VWF complexes.

Using ristocetin-induced platelet agglutinations, we show that plasmin cleavage of VWF does not impair its platelet-binding capacity, whereas plasmin-mediated cleavage of GPIbα reduces the ability of platelets released from agglutinates to bind VWF. Furthermore, platelets in suspension are relatively resistant to plasmin cleavage. We therefore propose that VWF binding may enhance GPIbα cleavage by recruiting plasmin(ogen) to the platelet surface.

In a TTP mouse model, plasminogen activation led to a VWF-dependent reduction in GPIbα detectability, although to a lesser extent than observed in vitro. In patients with acute TTP, soluble GPIbα levels were elevated, indicating increased GPIbα shedding during attacks of thrombotic microangiopathy, although the extent to which this is plasmin-mediated remains unclear.

Together, our findings demonstrate that plasmin cleavage of GPIbα drives the disruption of ristocetin-induced agglutinates, while its contribution to the breakdown of platelet-VWF complexes in vivo appears limited.

## Introduction

Von Willebrand factor (VWF) is a large multimeric glycoprotein that is essential for primary hemostasis by recruiting platelets to sites of vascular injury.^1^ VWF circulates in a globular conformation with its platelet-binding A1 domain shielded, whereas newly secreted or collagen-bound VWF is uncoiled and capable of engaging the glycoprotein (GP) Ibα receptor on platelets.^2,3^ The adhesive potential of unfolded VWF increases with multimer size. To prevent uncontrolled platelet binding, VWF multimer size is regulated by the metalloprotease ADAMTS13 (a disintegrin and metalloproteinase with a thrombospondin type 1 motif, member 13), which cleaves VWF in its A2 domain to prevent the accumulation of ultra-large, highly adhesive forms.^4,5^ When this regulation falls short, platelet-VWF complexes can occlude the (micro)circulation. This is most strikingly exemplified by immune-mediated thrombotic thrombocytopenic purpura (TTP), where autoantibodies inhibit ADAMTS13, leading to unpredictable attacks of widespread microvascular thrombosis.^6^

In previous work, we provided evidence of increased endogenous plasmin generation in experimental models of TTP and in TTP patients during acute attacks.^7,8^ Building on these findings, we proposed that plasmin may function as an auxiliary VWF-cleaving protease to clear microvascular occlusions. To enhance this process therapeutically, we developed the VWF-targeting plasminogen activator Microlyse, which promotes local plasmin formation at sites of VWF accumulation. We showed that Microlyse treatment attenuates thrombocytopenia and tissue injury in a murine TTP model, supporting the concept that targeted plasmin-mediated VWF proteolysis contributes to microthrombus resolution.^9^

We recently expanded on this concept by characterizing VWF cleavage during plasmin-mediated breakdown of platelet-VWF complexes. We found that plasmin cleavage results in the formation of a largely polymeric degradation product of VWF (cVWF). In patients with TTP, cVWF is formed during acute attacks, suggesting that it reflects ongoing microvascular occlusion. In a murine TTP model, we showed that VWF clearance following Microlyse treatment is accompanied by an increase in cVWF formation, suggesting that cVWF may also hold value as a monitoring tool for therapeutic plasminogen activation.^10^

These observations raised questions about the functional consequences of plasmin cleavage, as it is unclear whether the disruption of platelet-VWF interactions is solely attributable to VWF proteolysis or whether it also involves proteolysis of its platelet receptor GPIbα, which has been reported to be susceptible to plasmin cleavage.^11,12^ In this study, we therefore investigated how plasmin-mediated proteolysis of VWF and GPIbα contribute to the breakdown of platelet-VWF complexes.

## Materials and Methods

Additional methods and a full list of reagents are provided in the Supplementary Materials.

### Generation of plasmin-cleaved VWF (cVWF)

Lyophilized Haemate-P (plasma-derived factor VIII/VWF concentrate) was reconstituted in distilled water according to the manufacturer’s instructions. Plasminogen was purified from human citrated plasma as described previously^13^ and activated with streptokinase (2000 IU/mL) for 15 min at 37°C. Haemate-P (250 µg/mL) was then incubated with plasmin (143 μg/mL) at 37°C for 1 hour. The reaction was terminated by the addition of 200 μM of the small-molecular protease inhibitor PPACK, as previously validated.^10^

### Formation of agglutinates (platelet-VWF complexes)

Washed platelets were mixed with 0.55 μg/mL iloprost, 280 μM dRGDW and full-length VWF (Haemate-P) or cVWF as a source of VWF (18 μg/mL final concentration, unless indicated otherwise). 220 μL of this mix was pre-warmed for 5 minutes at 37°C in an aggregometer (CHRONO-LOG® Model 700, Chrono-log Corporation, Havertown, Pennsylvania, USA) and constantly mixed by a magnetic stirrer (900 RPM). After one minute, 12 μL ristocetin (for a final concentration of 776 μg/mL) was added to the platelet mix to induce agglutinate formation. Light transmission was recorded for indicated durations.

### Plasmin-mediated agglutinate breakdown and re-agglutination

Agglutinates were allowed to form for six minutes after ristocetin addition, after which 50 μL of plasmin (0, 37.5, 75 or 150 μg/mL final concentration) was added. After 30 minutes, plasmin was inactivated by the addition of PPACK (5 μL, 209 μM final concentration). For experiments where plasmin was inhibited by aprotinin, a final concentration of 150 KIU/mL was added after agglutinates had formed, 1 minute prior to plasmin addition. To fix the platelets for experiments where flow cytometry analysis was performed, a 5 μL sample was collected into 290 µL of fixative solution (137 mM NaCl, 2.7 mM KCl, 1.12 mM NaH_2_PO_4_·H_2_O, 1.15 mM KH_2_PO_4_, 10.2 mM Na_2_HPO_4_·2H_2_O, 4 mM EDTA, 1.11% formaldehyde, pH 6.8) after plasmin inactivation. A second dose of ristocetin (15 μL, for a final concentration of 758 μg/mL) was then added to assess re-agglutination capacity. Where indicated, samples were supplemented with full-length VWF (18 µg/mL final concentration) after plasmin inactivation, followed by ristocetin to assess re-agglutination capacity (761 μg/mL final concentration). For quantification of soluble GPIbα (sGPIbα) release by enzyme-linked immunosorbent assay (ELISA), samples were collected at the end of the aggregometry run and centrifuged twice for 15 minutes at 400xg to ensure complete removal of residual platelets.

### Plasmin-mediated GPIbα cleavage in washed platelet suspensions

Platelet suspensions were incubated with streptokinase-activated plasminogen (50 μL, final concentration 0, 37.5, 75 or 150 µg/mL) for 30 minutes at 37°C without exogenous VWF to specifically assess the effects of plasmin on platelets independently of its activity on VWF. Plasmin activity was inhibited by the addition of PPACK (5 μL, 218 μM final concentration). For flow cytometry, a 5 μL sample was taken after plasmin inactivation and collected into 290 µL of fixative solution. For sGPIbα measurements by ELISA, samples were collected at the end of the aggregometry run and spun down twice for 15 minutes at 400 xg to collect platelet-free supernatants. Haemate-P (12 μL, final concentration 18 μg/mL) was then added to the platelet mix as a source of VWF, followed by ristocetin (14 μL for a final concentration of 737 μg/mL) to trigger agglutinate formation.

### Time course analysis of plasmin-mediated GPIbα cleavage

Washed platelet suspensions were mixed with 0.55 μg/mL iloprost and 280 μM dRGDW. In a subset of samples, plasmin (75 µg/mL) was added directly and incubated for 0, 2.5, 5, 10, 20 or 30 minutes under continuous stirring in an aggregometer (900 RPM, 37°C). In a second subset of samples, VWF (18 µg/mL) and ristocetin (776 µg/mL) were first added to induce agglutination, which was allowed to proceed for six minutes before adding plasmin (75 µg/mL). After incubation for the indicated durations, PPACK (218 µM) was added to all conditions to quench plasmin activity. Samples were then centrifuged twice at 400 xg for 15 minutes to remove platelets, and sGPIbα levels were quantified by ELISA in the platelet-free supernatants.

### Flow cytometry of washed platelets

A 5 µL sample was collected into 290 µL of fixative solution and incubated for 15 minutes to preserve surface markers. Samples were then centrifuged for 15 minutes at 400 xg and supernatants were discarded. An antibody mixture for the detection of GPIbα (variable domain of heavy-chain-only antibody (V_H_H); clone ‘19’; in-house), VWF (polyclonal anti-VWF-Alexa Fluor 488; ab8822; Abcam), and GPIIb (monoclonal anti-CD41a-PerCP; clone MEM-06; ab134373; Abcam) was prepared in HT buffer (pH 7.4) and 50 µL was added to the platelet pellet, followed by 30 minutes incubation at room temperature (RT). Stained samples were fixed with 290 µL of fixative and incubated for 15 minutes, after which the fixed sample was further diluted two times in fixative. Samples were analyzed on a FACS Canto II using FACS Diva software version 8.0.1. Platelets were gated based on forward and sideward scatter and CD41+ events were determined in this population. Data acquisition aimed for 10,000 events per sample and the median fluorescence intensity (MFI) of a minimum of 8,000 gated platelets was determined per sample.

### Soluble GPIbα (sGPIbα) ELISA

sGPIbα-specific capture antibody clone ‘24B3’ ^14^ was coated onto 96-well MaxiSorp plates (5 µg/mL in PBS, 50 µL per well) overnight at 4°C. Plates were rinsed 3 times with 0.01% (v/v) Tween-20 in PBS (PBS-T). Plates were then blocked with 1% (w/v) skim milk in PBS (block buffer; 200 µL per well) for 1 hour at RT while shaking (600 RPM). Platelet-free supernatants were diluted 8 times in block buffer and normal pool plasma was serially diluted in block buffer and used as an assay standard, based on a sGPIbα plasma concentration of 2 µg/mL.^15^ Samples were applied and incubated at RT for 1 hour while shaking, after which plates were rinsed 3 times with PBS-T. Bound sGPIbα was detected with biotinylated sGPIbα-specific detection antibody clone ‘6B4’ ^16^ (1 µg/mL; 50 µL per well). Plates were rinsed 3 times after incubation with detection antibody for 1 hour at RT, after which an additional incubation was performed with poly-horseradish peroxidase–labeled streptavidin (diluted 4000 times in block buffer; 50 μL per well) for 1 hour at RT while shaking. Plates were rinsed 3 times with PBS-T and developed by the addition of 100 μL of 3,3′,5,5′-tetramethylbenzidine substrate. Substrate development was allowed for a maximum of 30 minutes, after which 50 μL of H_2_SO_4_ (0.3 M) was added for end point absorption measurements at 450 nm. Results were analyzed by GraphPad Prism 10.4.0 using a sigmoidal 4PL fit model to which sample concentrations of sGPIbα were related.

### Plasminogen binding studies

An antibody mixture for the detection of endogenous VWF and plasminogen was prepared in HT buffer (pH 7.4), containing polyclonal anti-VWF-Alexa Fluor 488 (ab8822; Abcam) and mouse monoclonal anti-plasminogen (clone 10A1, Invitrogen, LF-MA0170). A competing GPIbα-targeting V_H_H (clone ‘17’; in-house; 75 or 750 nM final concentration) previously described by Sanrattana et al.,^17^ ε-aminocaproic acid (εACA; 10 mM), or buffer as control was then added to the antibody mix. Ristocetin (750 µg/mL final concentration) was added to induce VWF binding to platelets where indicated, and buffer was used as a control.

Separately, citrated whole blood was incubated for 10 minutes at RT with iloprost (600 ng/mL), εACA (10 mM) and/or the competing GPIbα-targeting V_H_H (75 or 750 nM final concentration), depending on the condition. These reagents were pre-incubated with whole blood to ensure that GPIbα and lysine-binding sites were blocked prior to VWF activation by ristocetin, thus preventing premature binding of VWF and plasminogen. After incubation, 5 µL of whole blood was added to 50 µL of the antibody mix and incubated for 20 minutes at RT. Samples were then incubated in fixative for 10 minutes and centrifuged at 340 xg for 15 minutes. Supernatants were discarded, and pellets were resuspended in 50 µL of secondary goat anti-mouse IgG1-PE antibody (SouthernBiotech) in fixative and incubated for 20 minutes at RT. Finally, samples were diluted 1:20 in fixative and analyzed by flow cytometry. Platelets were gated based on forward and side scatter properties.

### In vivo studies

All animal studies were performed at the animal institution of KU Leuven in accordance with protocols approved by the Institutional Animal Care and Use Committee (project number: P101/2024). Female and male *Adamts13*^−/−^ mice (CASA/Rk-C57BL/6J-129X1/SvJ background) of 8-12 weeks old were used. Mice were anesthetized using isoflurane and intravenously injected with 500 U/kg recombinant human VWF (Veyvondi®) mixed with 20 µg/g human plasminogen or an equal volume of saline. After 5 minutes, mice received an intravenous injection with 1 U/g streptokinase or saline.^7^ Blood was collected 24 hours after the injections in citrate (5:1 vol/vol of blood:3.8% trisodium citrate) and EDTA (10:1 vol/vol of blood:0.5 M EDTA) for further analysis. Total blood cell counts were analyzed on EDTA blood samples using the Vetscan HM5 (Zoetis, Parsippany, New Jersey, USA). Citrated whole blood was incubated for 10 minutes with antibodies for the detection of GPIbα (CD42b, Emfret), GPIIb (CD41, BioLegend) and VWF (Emfret). Fluorescence was detected using flow cytometry (Cytoflex S, Beckman Coulter, Brea, California, USA). Platelets were gated based on their forward and sideward scatter properties, and CD41^+^ events were identified within this population. The median fluorescence intensity of 10,000 gated platelets was determined.

### Human plasma studies

Citrated plasma from acute iTTP patients (n=83) was collected at the Department of Blood Transfusion Medicine at Nara Medical University, under the approval of the Ethics Committee of the Nara Medical University and conducted under the tenets of the Declaration of Helsinki. Samples were taken during the acute phase, prior to initial treatment. Plasma from healthy volunteers (n=43) was prepared from blood samples taken by venous phlebotomy with ethical approval of the University Medical Center Utrecht. Plasma samples were diluted 32 times in block buffer for evaluation of sGPIbα levels by ELISA. Plasma levels of cVWF were measured by ELISA using variable domain of heavy-chain-only antibodies as previously described by El Otmani et al.^19^

### Data analysis

Statistical analyses were performed using one-way analysis of variance (ANOVA) or two-way ANOVA with Tukey’s post-hoc test for multiple comparisons. For measurements of sGPIbα in iTTP patients, normality was assessed using D’Agostino & Pearson, Anderson-Darling, Shapiro-Wilk, and Kolmogorov-Smirnov tests, followed by the Mann Whitney test for comparisons with healthy individuals. Correlations were evaluated using Spearman’s rank correlation coefficient (r). *P* < 0.05 was considered statistically significant. All data analyses were performed using Prism version 10.4.0 (GraphPad Software, LLC).

## Results

### Plasmin-mediated breakdown of ristocetin-induced platelet agglutinates cannot be explained by VWF cleavage alone

To investigate whether proteolysis by plasmin affects VWF function, we first evaluated whether plasmin-cleaved VWF (cVWF) can still support the formation of platelet-VWF complexes. cVWF was generated by incubating full-length plasma-derived VWF with plasmin as described previously by El Otmani et al. (Supplemental Figure 1).^10^ Platelet-VWF complexes were modeled in vitro using ristocetin-induced platelet agglutination.

No spontaneous agglutination occurred in the absence of ristocetin (Figure 1A). At 18 µg/mL, cVWF supported ristocetin-induced agglutination to a similar extent as full-length VWF. Similar agglutination was observed at 9 µg/mL for both cVWF and full-length VWF, while approximately 5% more platelets remained in suspension at 4.5 µg/mL cVWF compared to full-length VWF.

**Figure 1.**
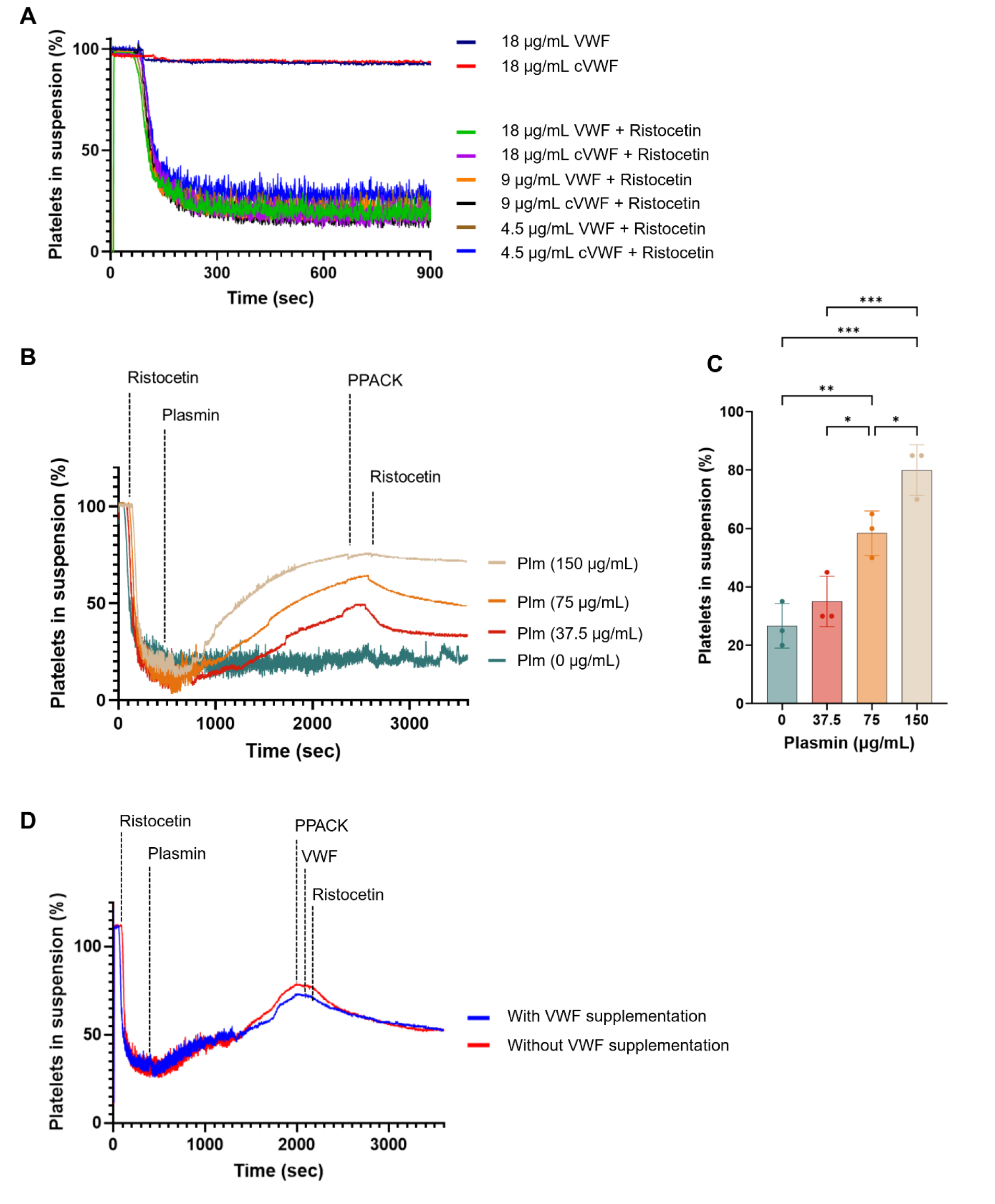
Agglutination capacity following plasmin exposure. (A) Washed platelets were mixed with full-length VWF or cVWF at indicated concentrations. Ristocetin was added to induce agglutination. (B) Washed platelets mixed with VWF were stimulated with ristocetin to form agglutinates. Plasmin was incubated for 30 minutes at indicated concentrations and inactivated with PPACK. A second ristocetin dose was added to assess re-agglutination capacity. (C) Percentage of platelets remaining in suspension after the second ristocetin trigger. (D) Agglutination after supplementation with fresh full-length VWF to agglutinates treated with plasmin (75 μg/mL). Data represent three independently executed experiments. Scatter plots with bar graphs show means ±SD. Data were analyzed by one-way ANOVA followed by Tukey’s multiple comparisons test. * *P* < 0.05; ** *P* < 0.01; *** *P* < 0.001. Plm = plasmin.

Next, we assessed the effect of plasmin on preformed platelet-VWF complexes. Plasmin treatment for 30 minutes caused a dose-dependent breakdown of agglutinates (Figure 1B), which was prevented by aprotinin, confirming plasmin dependency (Supplemental Figure 2A). Western blot analysis confirmed VWF cleavage under these conditions (Supplemental Figure 2B, C). After quenching plasmin activity with PPACK, a second dose of ristocetin was added to assess the remaining agglutination capacity (Figure 1B). The capacity to re-form agglutinates decreased progressively with increasing plasmin concentrations, as reflected by the higher percentage of platelets remaining in suspension (Figure 1B, C). As a control, we confirmed that doubling the ristocetin dose did not induce spontaneous agglutinate formation in the absence of VWF (Supplemental Figure 3). To test whether the reduced re-agglutination was due to impaired VWF function, we added fresh full-length VWF to plasmin-treated samples following plasmin inactivation (Figure 1D). Ristocetin was then added to assess whether agglutination could be restored. However, agglutination did not improve, indicating that the functional impairment was not due to VWF degradation alone.

### Plasmin induces GPIbα shedding, which is amplified by VWF binding

Since VWF cleavage alone did not account for the observed loss of agglutination, we next examined how plasmin affects surface levels of GPIbα. Platelet suspensions were incubated with plasmin for 30 minutes after which GPIbα surface levels were measured by flow cytometry. GPIbα signal progressively declined with increasing plasmin concentrations, reaching a maximal reduction of ∼33% in median fluorescence intensity (MFI) relative to baseline (untreated control) (Figure 2A). In parallel, we analyzed GPIbα detectability on platelets released by plasmin from ristocetin-induced agglutinates, as described above (Figure 1B). A control without plasmin could not be included here, as most platelets remained in complex with VWF, limiting the recovery of single, unbound platelets for reliable analysis (Supplemental Figure 4). Under these conditions, the decrease in GPIbα detectability was significantly greater than in platelet suspensions, with a maximal reduction in MFI of ∼75% relative to baseline.

**Figure 2.**
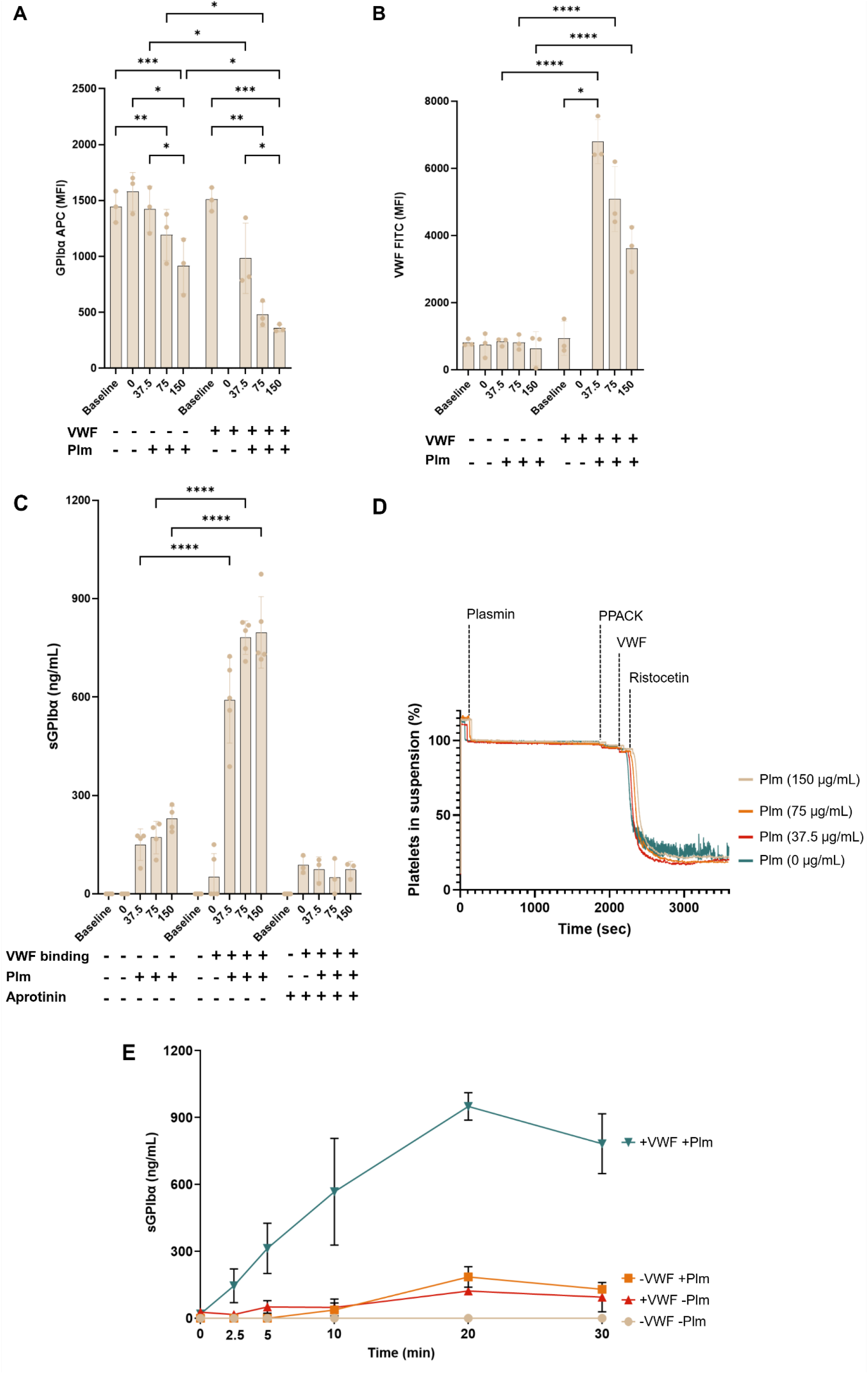
Plasmin cleaves GPIbα during agglutinate breakdown. (A, B) GPIbα and VWF detectability was assessed by flow cytometry after 30 minutes of plasmin incubation at indicated concentrations. (C) sGPIbα levels were measured by ELISA in platelet-free supernatants from samples treated with indicated plasmin concentrations in the absence or presence of VWF and/or aprotinin (150 KIU/mL). (D) Washed platelet suspensions were pre-incubated with indicated concentrations of plasmin for 30 minutes in the absence of exogenous VWF. After plasmin inactivation with PPACK, full-length VWF and ristocetin were added to assess agglutination. (E) Time course analysis of sGPIbα release upon plasmin (75 μg/mL) treatment, measured by ELISA in platelet-free supernatants. Data represent three independently executed experiments. Scatter plots with bar graphs and XY plots show means ±SD. Data were analyzed by two-way ANOVA followed by Tukey’s multiple comparisons test. * *P* < 0.05; ** *P* < 0.01; *** *P* < 0.001; **** *P* < 0.0001. MFI = median fluorescence intensity. Plm = plasmin. Baseline= untreated control. 0 = incubation without plasmin.

Only minimal platelet-bound VWF was detected in platelet suspensions incubated without exogenous VWF (Figure 2B). Platelets released from agglutinates displayed higher VWF levels due to ristocetin-induced VWF binding, which declined progressively with increasing plasmin concentrations but did not reach statistical significance. As platelets were incubated in the presence of inhibitors of platelet activation, P-selectin was undetectable under all conditions (data not shown).

To determine whether the reductions in GPIbα detectability reflected proteolytic shedding, soluble GPIbα (sGPIbα) levels were measured in platelet-free supernatants by ELISA (Figure 2C). At baseline, sGPIbα was undetectable. Plasmin treatment of platelet suspensions led to sGPIbα release, with levels reaching up to ∼230 ng/mL. In supernatants from ristocetin-induced agglutinates, sGPIbα (∼50 ng/mL) was detected even without plasmin. The addition of plasmin further increased sGPIbα, reaching a plateau at ∼800 ng/mL. Notably, the difference in sGPIbα levels between platelet suspensions and agglutinates was larger than the difference observed by flow cytometry. When plasmin activity on agglutinates was inhibited by co-incubation with aprotinin, sGPIbα levels remained comparable to those observed without plasmin.

Next, we examined whether the GPIbα loss in plasmin-treated platelet suspensions (Figure 2A) impaired their ability to agglutinate when full-length VWF and ristocetin were added after plasmin inactivation (Figure 2D). In contrast to the impaired agglutination observed when platelets were treated with plasmin after ristocetin-induced VWF binding (Figure 1B), agglutination remained similar to the control without plasmin.

To further examine the kinetics of plasmin-mediated GPIbα cleavage, we performed a time course analysis in which platelets were treated with plasmin (75 µg/mL) either without exogenous VWF or with ristocetin-induced VWF binding (Figure 2E). Without VWF, plasmin induced a gradual increase in sGPIbα over time, whereas when VWF was bound, plasmin markedly accelerated and amplified sGPIbα release, which approached a plateau after 20 minutes.

### VWF facilitates plasminogen binding to platelets

Since binding of VWF to GPIbα amplifies plasmin-mediated GPIbα shedding, we hypothesized that platelet-bound VWF might act as a scaffold that directly binds plasminogen and recruits it to the platelet surface. To investigate this, we performed flow cytometry analysis of endogenous VWF and plasminogen binding to platelets in citrated whole blood. Ristocetin was added to induce VWF binding to GPIbα. To block this interaction, a GPIbα-targeting variable domain of heavy-chain-only antibody^17^ (V_H_H) was added at a low (75 nM) and a high (750 nM) concentration. The lysine analog ε-aminocaproic acid (εACA) was used to inhibit the lysine-dependent binding of plasminogen to the A1 domain of VWF.^7^

Ristocetin significantly increased VWF binding (Figure 3A). The blocking V_H_H dose-dependently reduced this interaction, while the addition of εACA did not affect VWF binding. Next, we assessed plasminogen binding (Figure 3B). Without ristocetin, plasminogen binding was low, but it increased significantly upon ristocetin-induced VWF binding. This increase was significantly reduced by εACA, and further suppressed by the blocking V_H_H in a dose-dependent manner, suggesting that plasminogen binding involves lysine-dependent binding to VWF and is facilitated by VWF-GPIbα interactions.

**Figure 3.**
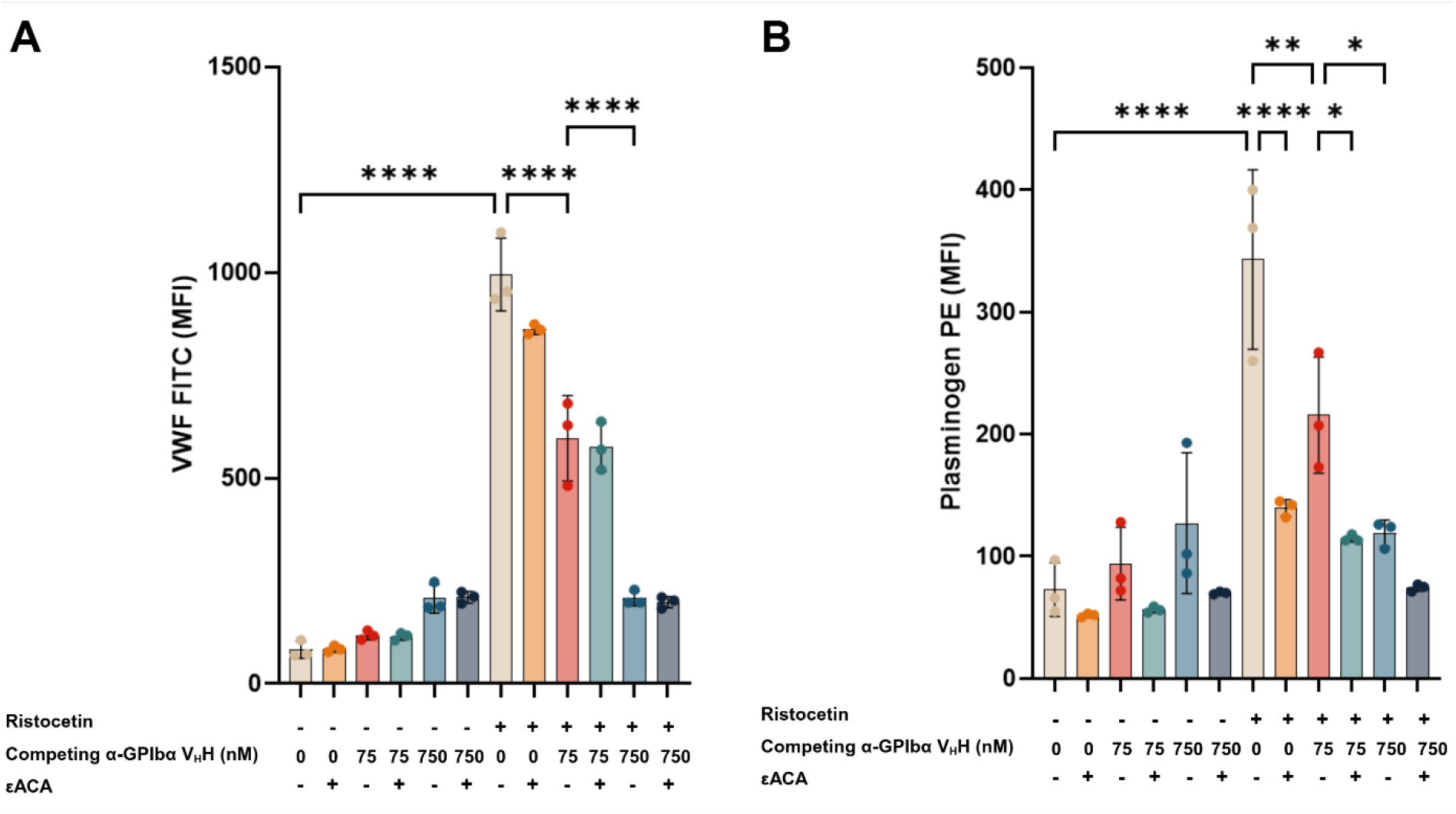
VWF facilitates plasminogen binding. (A) VWF binding to platelets in citrated whole blood was assessed by flow cytometry in the presence or absence of ristocetin, a blocking GPIbα-targeting V_H_H (75 or 750 nM) and εACA (10 mM). (B) Plasminogen binding was measured under the same conditions. Data represent three independent experiments and are displayed as scatter plots with bar graphs showing means ±SD. Data were analyzed by two-way ANOVA followed by Tukey’s multiple comparisons test. Key comparisons are shown. * *P* < 0.05; ** *P* < 0.01; **** *P* < 0.0001. MFI = median fluorescence intensity.

### VWF enhances plasmin-mediated loss of GPIbα detectability in vivo

Building on our in vitro findings that VWF binding increases plasmin-mediated cleavage of GPIbα, we next explored whether this increase also occurs in vivo. For this purpose, we used ADAMTS13-deficient mice and a VWF-driven model of thrombotic microangiopathy that is used to study TTP.^7,9,10^ Platelet-VWF complexes were induced by administration of recombinant human VWF (rVWF), which was co-administered with human plasminogen (Plg). Five minutes later, streptokinase (Strep) was given to activate plasminogen. Additional groups received either rVWF alone, plasminogen and streptokinase without rVWF, or saline as a control. Whole blood was collected 24 hours after treatment for platelet count measurements and flow cytometry analysis.

rVWF administration resulted in a significant reduction in platelet counts (Figure 4A). In mice that received Plg/Strep treatment following rVWF administration, mean platelet counts recovered partially to ∼75% of those in saline-treated mice. Platelet-bound VWF levels were significantly higher in mice that received rVWF alone than in saline-treated mice, but did not differ from the group that received rVWF and Plg/Strep (Figure 4B). GPIbα detectability was significantly lower in mice that received both rVWF and Plg/Strep than in those treated with Plg/Strep alone, with a mean MFI reduction of ∼7.7% (Figure 4C). This reduction is approximately tenfold lower than the maximum decrease in GPIbα detectability observed in vitro, where we measured a reduction of up to ∼75% (Figure 2A).

**Figure 4.**
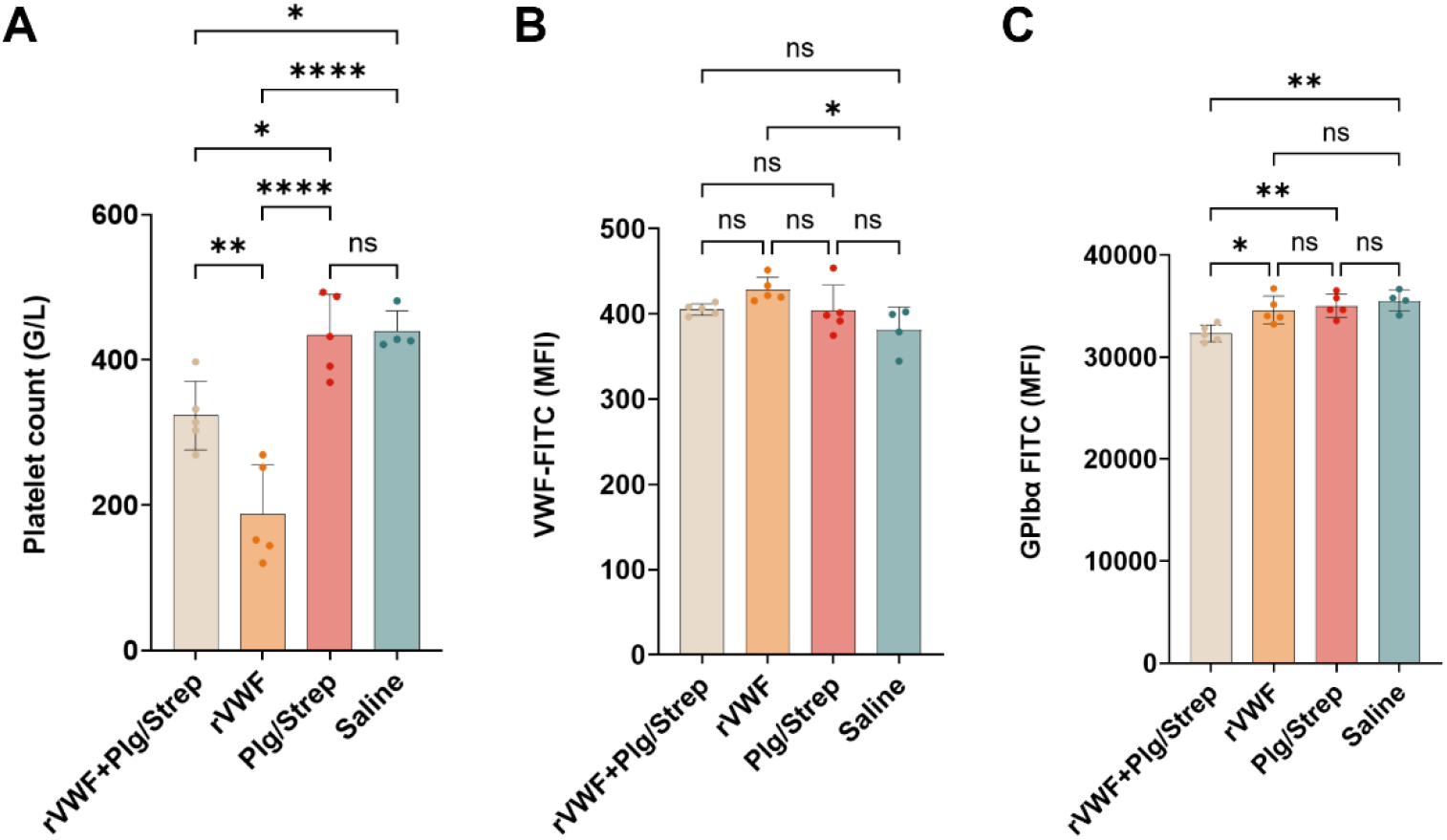
Plasmin-mediated modulation of GPIbα in vivo. *Adamts13*^−/−^ mice were treated with either saline, recombinant human VWF (rVWF), rVWF and plasminogen followed by streptokinase (rVWF + Plg/Strep), or plasminogen followed by streptokinase alone (Plg/Strep). Whole blood was collected 24 hours after treatment for (A) platelet count measurements and (B,C) flow cytometry analysis of platelet-bound VWF and GPIbα. Data are presented as scatter plots with bar graphs showing means ±SD. Data were analyzed by one-way ANOVA. ns, nonsignificant; \**P* < 0.05; \*\**P* < 0.01; \*\*\**P*<0.001; \*\*\*\**P* < 0.0001.

### sGPIbα levels are elevated during acute attacks in patients with TTP

Acute TTP is characterized by platelet- and VWF-rich microthrombi. As we showed in previous studies, this coincides with increased plasmin generation and elevated plasma levels of cVWF.^7,8,10^ Following our in vitro observations that plasmin more readily acts on GPIbα when it is engaged in platelet-VWF complexes, we next asked whether GPIbα is similarly targeted during TTP attacks, where such complexes are abundant. To test this, we measured sGPIbα and cVWF levels in plasma samples from immune-mediated TTP patients during acute episodes (n=83) and compared them to plasma obtained from healthy individuals (n=43). Patient characteristics are listed in Table 1.

**Table 1.** Patient and healthy donor characteristics.

|  | N | Sex (female/male) | Age (y), median (IQR) |
| --- | --- | --- | --- |
| Acute TTP | 83 | 48/35 | 58 (41.5, 69.0) |
| Normal healthy donors | 43 | 23/20 | 42 (32.0, 55.0) |

sGPIbα levels were significantly higher in TTP patients than in healthy individuals, with median levels of 3677 and 1544 ng/mL, respectively (Figure 5A). A weak correlation was observed between sGPIbα and cVWF concentrations (r = 0.31; *P* < 0.01) (Figure 5B).

**Figure 5.**
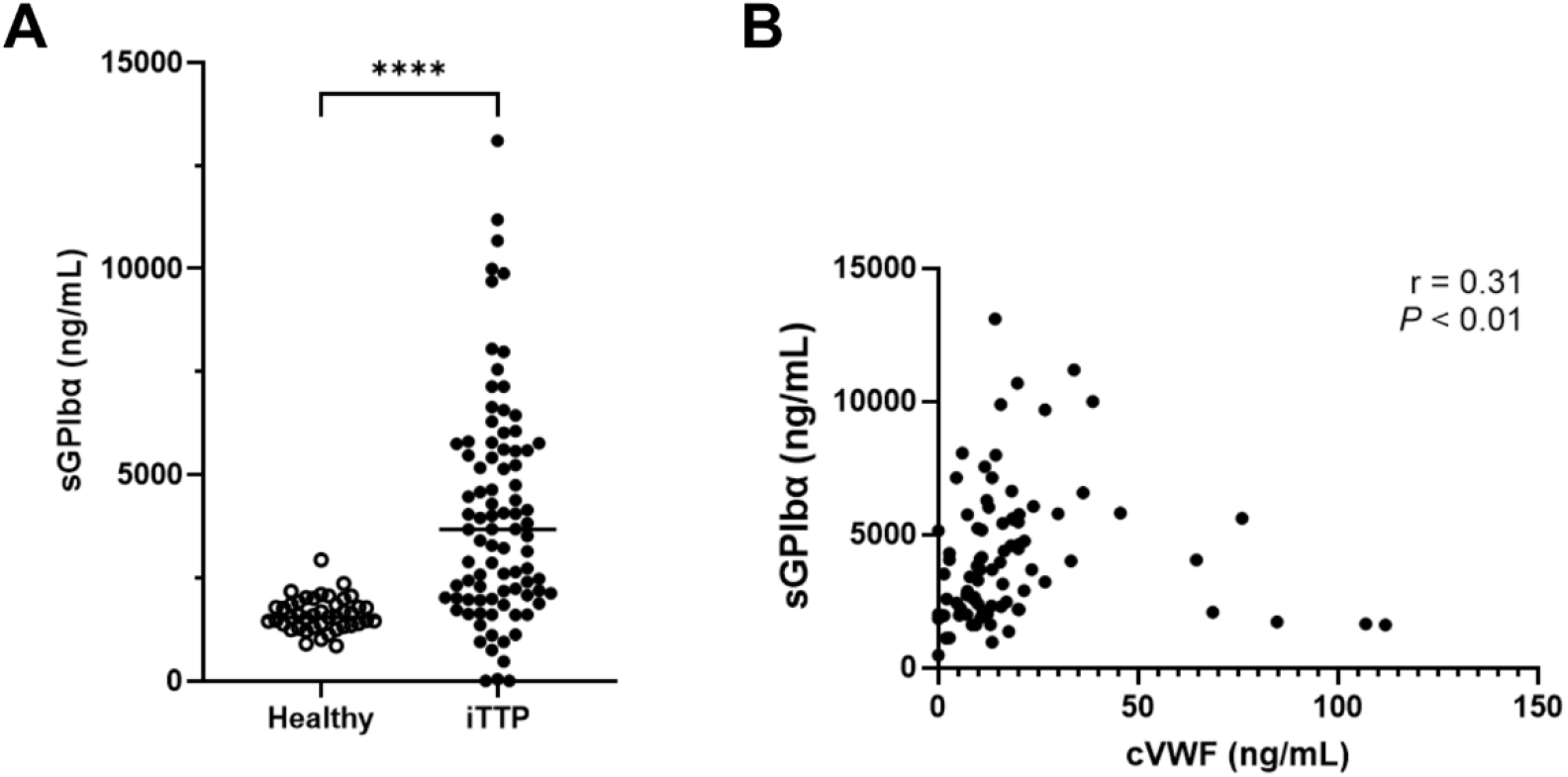
sGPIbα levels are elevated in TTP patients during attack. (A) Plasma sGPIbα levels in TTP patients experiencing an acute attack (n=83) compared to healthy controls (n=43). Data are shown as scatter plots with medians and were analyzed by Mann-Whitney test. \*\*\*\**P* < 0.0001. (B) Correlation between plasma sGPIbα and cVWF levels during acute TTP evaluated by Spearman’s rank correlation coefficient (r).

## Discussion

We here show that cVWF retains the ability to support ristocetin-induced platelet agglutination, indicating that plasmin-mediated VWF cleavage alone does not account for the breakdown of platelet-VWF complexes in vitro. Instead, we found that plasmin-mediated shedding of GPIbα reduces the capacity of platelets released from agglutinates to reform agglutinates, whereas platelets in suspension are relatively resistant to plasmin cleavage. This functional impairment could not be restored by the addition of full-length VWF after plasmin treatment, indicating that loss of GPIbα, rather than VWF proteolysis, drives agglutinate breakdown in ristocetin-driven experiments. Our finding that cleavage of GPIbα is enhanced by VWF binding supports a model in which VWF recruits plasmin(ogen) to the platelet surface and brings it in a position to cleave GPIbα.

Plasminogen binding increased when VWF was bound to GPIbα, which could be blocked by competition for lysine binding sites by εACA or by V_H_H-mediated displacement of VWF from GPIbα. This suggests that VWF might serve as a scaffold for plasmin(ogen), likely via a lysine-rich patch in its A1 domain,^7^ thereby promoting localized proteolysis of GPIbα. Moreover, VWF binding has been reported to promote clustering of GPIbα,^18^ which could increase the local density of GPIbα available for proteolysis, potentially contributing to enhanced shedding.

VWF binding under shear stress exerts mechanical forces that induce conformational changes in GPIbα, including unfolding of the mechanosensitive domain (MSD),^19,20^ thereby facilitating access of proteases to otherwise shielded regions. Although the plasmin cleavage site in GPIbα has not yet been identified, VWF-induced conformational changes could potentially enhance plasmin-mediated proteolysis of the receptor. Notably, low-level shedding following VWF binding was also observed in the absence of plasmin, suggesting involvement of endogenous platelet proteases. One possible contributor is a disintegrin and metalloproteinase (ADAM) 17, the main catalyst of GPIbα shedding, which cleaves the receptor within the MSD. However, ADAM17-mediated proteolysis typically proceeds with slow kinetics, requires sustained platelet activation and was recently found to occur mainly intracellularly.^21^ As platelet activation was pharmacologically inhibited in all in vitro experiments, ADAM17 is therefore an unlikely mediator of the observed proteolysis.

Plg-R_KT_ has been described as a lysine-dependent plasminogen receptor that is localized intracellularly and requires platelet activation to translocate to the platelet surface, where it retains platelet-derived plasminogen.^22^ Our data show that plasminogen binding occurs independently of platelet activation, suggesting that Plg-R_KT_ is unlikely to account for the observed binding. Experiments with activated platelets that involve V_H_H-mediated blockade of the VWF-GPIbα interaction or direct inhibition of Plg-R_KT_ could help clarify its potential contribution. It should be noted, however, that in acute TTP, minimally activated platelets form complexes with VWF, yet both endogenous plasmin and Microlyse act on these complexes. This argues against a dominant role of Plg-R_KT_ in mediating plasminogen recruitment under these conditions.

Our findings suggest that the mechanism of action of VWF-targeting thrombolytics like Microlyse may extend beyond VWF degradation and could also involve GPIbα cleavage. This could be beneficial in preventing the reformation of platelet-VWF complexes, as our in vitro results show that a loss of surface GPIbα is associated with a reduction in agglutination capacity. Based on the observation that cVWF can still support ristocetin-induced agglutination, GPIbα cleavage could prevent platelet re-adhesion after VWF proteolysis, although it is uncertain to what extent cVWF retains platelet-binding capacity in vivo. Another implication may be that because platelet suspensions are relatively resistant to plasmin cleavage, circulating platelets not participating in thrombi may similarly remain largely unaffected by plasmin, which would represent an important safety advantage.

On the other hand, platelets liberated from occlusions by plasmin cleavage may be functionally compromised when they re-enter the circulation. Defining the threshold of GPIbα loss at which this functional impairment occurs may aid in stratifying bleeding risk following therapeutic plasminogen activation. It should be noted that Microlyse administration in a TTP mouse model did not lead to a prolongation of bleeding time.^9^ Furthermore, it has been reported that platelets can replenish their surface GPIbα pool by recruitment of intraplatelet GPIbα, suggesting that any potential loss of function may be transient.^23^

A VWF-dependent reduction in GPIbα detectability was also observed in a mouse model of TTP following plasmin treatment, but this effect was markedly lower than observed in vitro. This highlights important differences between our experimental models, including the absence of shear forces and of natural inhibitors of fibrinolytic activity in vitro. Moreover, ristocetin-driven agglutination does not fully recapitulate physiological VWF-GPIbα interactions, as it is an artificial trigger for VWF unfolding that can also induce interactions with other proteins aside from VWF, including GPIbα.^24,25^ Studies using shear-unfolded VWF secreted from endothelial cells under flow could offer insight into how plasmin-mediated GPIbα proceeds under more physiological conditions.

The smaller effect in vivo may also result from the rapid clearance of platelets that have lost GPIbα; given the five day lifespan of murine platelets, about 20% of circulating platelets are newly formed every 24 hours, which could lead to an underestimation of the true extent of GPIbα loss when measured after 24 hours.^20,26,27^ Assessments of earlier time points together with a time course analysis of murine sGPIbα are therefore needed to better understand the dynamics of this process. These studies should also examine the impact of GPIbα loss on platelet agglutination capacity.

Another consideration is the extent to which murine GPIbα is susceptible to cleavage by human plasmin. Mice are much less sensitive to thrombolytic therapies than humans and require doses about tenfold higher than the established clinical dose,^28^ which highlights interspecies differences in the fibrinolytic system that could also affect cleavage of murine substrates by human plasmin. In addition, murine GPIbα binds poorly to human VWF,^29^ which may limit VWF-dependent plasmin cleavage of GPIbα in the present model. This should also be taken into account when interpreting the hemostatic outcomes following Microlyse treatment in this model. Species-matched plasminogen-activators and experimental models resistant to plasmin-mediated proteolysis of either GPIbα or VWF could be valuable tools to determine how each plasmin target contributes to the breakdown of platelet-VWF complexes in vivo.

In patients with acute TTP, we previously demonstrated that attacks are accompanied by increased plasmin activity and elevated cVWF levels, which correlate with each other and with thrombocytopenia.^10^ The increased sGPIbα levels observed during acute TTP could therefore, at least in part, reflect plasmin-mediated GPIbα proteolysis. However, circulating sGPIbα likely consists of a heterogeneous pool generated by different enzymes. Indeed, other proteases, including calpain, have also been implicated in GPIbα cleavage during TTP attacks.^30^ Therefore, the extent to which this rise in sGPIbα is actually plasmin-dependent is unclear. Although the weak correlation between sGPIbα and cVWF suggests a possible link with plasmin activity, assessing markers of fibrinolytic activity such as plasmin-α2-antiplasmin complexes could provide further insight into the involvement of plasmin in this process.

In summary, our findings show that plasmin mediates the breakdown of platelet-VWF complexes not only through VWF cleavage but also through proteolytic shedding of GPIbα, which is amplified by VWF binding. This mechanism is most evident under ristocetin-induced conditions. While its contribution in vivo appears limited and difficult to interpret based on the current approaches, it warrants further investigation in more suitable models.

## Acknowledgements

The authors gratefully acknowledge Raymond Schiffelers, Sander Kooijmans, Karen Vanhoorelbeke, Simon De Meyer, Heyu Ni, and Guangheng Zhu for valuable discussions and input.

## Authors’ contributions

H.E.O. initiated and designed the study; R.F., E.I.M., A.H., C.T., and H.E.O. performed the experiments; K.S. and M.M. recruited the patients; R.F., E.I.M., C.T., and H.E.O. analyzed the data and interpreted the results. R.F. and H.E.O. wrote the manuscript with input from the other authors. All authors approved the final version of the manuscript.

## Conflict of interest disclosure

K.S. received speaker fees from Sanofi and participated in the advisory boards of Takeda and Alexion Pharma. M.M. provided consultancy services for Takeda, Alexion Pharma, and Sanofi; received speaker fees from Takeda, Alexion Pharma, Asahi Kasei Pharma, and Sanofi; and received research funding from Alexion Pharma, Chugai Pharmaceutical, Asahi Kasei Pharma, and Sanofi. The remaining authors declare no competing financial interests.

