## Supplementary Materials for "Plasmin-mediated cleavage of GPIbα contributes to breakdown of platelet-von Willebrand factor complexes"

### Supplementary Materials and Methods

#### Reagents

Tween-20, bovine serum albumin (BSA), glycine, sodium dodecyl sulfate (SDS), ε-Aminocaproic acid, aprotinin, bromophenol blue and DL-Dithiothreitol (DTT) were obtained from Sigma Aldrich, Saint Louis, Missouri, USA. 4-12% Bis-Tris Page gel, MOPS buffer and methanol were from Thermo Fisher, Waltham, Massachusetts, USA. Glucose and 4-(2-hydroxyethyl)-1-piperazineethanesulfonic acid (HEPES) were from VWR Avantor Life Science, Radnor, Pennsylvania, USA. $H_2SO_4$, ethanol and the Immobilon-FL membrane were obtained from Merck, Darmstadt, Germany. Tris was from Roche, Basel, Switzerland. Maxisorp plates were from Nunc, Thermo Fisher, Waltham, Massachusetts, USA. Haemate-P was from CSL Behring, King of Prussia, Pennsylvania, USA. Prostacyclin ($PGI_2$) was from Sanbio, Uden, the Netherlands. FPR-Chloromethylketone (PPACK) was from Enzyme Research Laboratories, South Bend, Indiana, USA. 3,3′,5,5′-Tetramethylbenzidine (TMB) was from Tebubio, Heerhugowaard, Netherlands. RGDW peptide was synthesized by NKI, Amsterdam, the Netherlands. Iloprost was from Bayer, Leverkusen, Germany. Ristocetin was from Biopool International, Ventura, California, USA. Streptokinase was from CSL-Behring, King of Prussia, Pennsylvania, USA. Odyssey blocking reagent was from LI-COR Biosciences, Lincoln, Nebraska, USA. Veyvondi ® was from Takeda, Tokyo, Japan.
Antibodies for western blot: rabbit polyclonal anti-human VWF (A0082, Agilent Dako, Santa Clara, California, USA). IRDye 680LT goat anti-rabbit IgG (P/N 926-68021, LI-COR Biosciences, Lincoln, Nebraska, USA). Antibodies for ELISA: mouse monoclonal capture antibody 24B3 and complementary biotinylated mouse monoclonal detection antibody 6B4 against human GPIbα were kindly provided by Karen Vanhoorelbeke (Leuven, Belgium). Capture $V_HH$ clone “G5” and biotinylated detection $V_HH$ clone “B9” against cVWF were produced in-house at UMC Utrecht (as described by El Otmani et al., Blood 2024). Poly-HRP-conjugated streptavidin (M2051, Streptavidin poly HRP, Sanquin, Amsterdam, Netherlands).
Antibodies for flow cytometry (in vitro studies): polyclonal anti-VWF-Alexa Fluor 488 antibody (ab8822) and mouse monoclonal anti-CD41a (GPIIb)-PerCP (clone MEM-06; ab134373) were from Abcam (Cambridge, United Kingdom). Mouse monoclonal anti-plasminogen (clone 10A1, LFMA0170) was from Fisher Scientific (Hampton, New Hampshire, USA). Goat $F(ab')_2$ Anti-Mouse IgG1-PE was from SouthernBiotech (Birmingham, AL, USA). For GPIbα detection, anti-GPIbα $V_HH$ clone ‘19’ -Alexa Fluor 647 was used. For competition assays, anti-GPIbα $V_HH$ clone ‘17’ was used, which was previously described (Sanrattana et al., J Thromb Haemost 2022). Antibodies for flow cytometry (in vivo studies): anti-CD42b-FITC and anti-VWF-FITC were from Emfret (Eibelstadt, Germany). Anti-CD41-Pacific Blue was from BioLegend (San Diego, California, USA).

**Blood collection and washed platelet preparation**

Whole blood from healthy donors was obtained under approval by the Medical Ethical Committee of the University Medical Center Utrecht. Blood samples were collected in sodium citrated collection tubes on the day of experiments and centrifuged at 160 xg for 15 minutes to obtain platelet-rich plasma (PRP). Platelet counts were determined by CELL-DYN Emerald Hematology Analyzer (Abbott Laboratories, Chicago, Illinois, USA). Prior to further centrifugation at 400 xg for 15 minutes for the separation of platelet-poor plasma (PPP), citrate-dextrose solution (ACD; 8.5 mM tri-sodium citrate, 7.1 mM citric acid, 11.1 mM D-glucose) was added to the undiluted PRP. PPP was removed and pellets were resuspended back to their initial volume using HEPES Tyrode buffer (HT buffer; 145 mM NaCl, 5 mM KCl, 0.5 mM $Na_2HPO_4$, 1 mM $MgSO_4$, 10 mM HEPES, 5.55 mM D-glucose, pH 6.5). $PGI_2$ (10 ng/mL) was added to platelet suspensions prior to another centrifugation step at 400 xg for 15 minutes. Supernatants were removed and the pellets were resuspended in HT buffer (pH 7.3); platelet counts were adjusted to $200 \times 10^9$/L.

**SDS PAGE and Western Blotting**

Samples were diluted in 3x sample buffer (30% glycerol, 0.18 M Tris-HCl, 6% SDS, Bromophenol blue, with or without 25 mM DTT for reduction), after which they were heated for 10 min at 95°C. Samples were separated on 4%-12% Bis-Tris gels at 165 V for 70 min in MOPS buffer and transferred to Immobilon-FL membranes at 125 V for 90 min in blotting buffer (25 mM Tris, 192 mM glycine, 20% (v/v) ethanol). Membranes were blocked for 2 h at room temperature (RT) with blocking buffer (Odyssey blocking reagent diluted 1:1 in TBS). Target proteins were detected by overnight incubation (at 4°C) of membranes. Membranes were washed with 0.1% (v/v) Tween in TBS (TBS-T) and incubated with secondary antibodies for 1 h at RT. Membranes were washed with TBS-T and analyzed on an Odyssey M scanner (LI-COR Biosciences, Lincoln, Nebraska, USA).

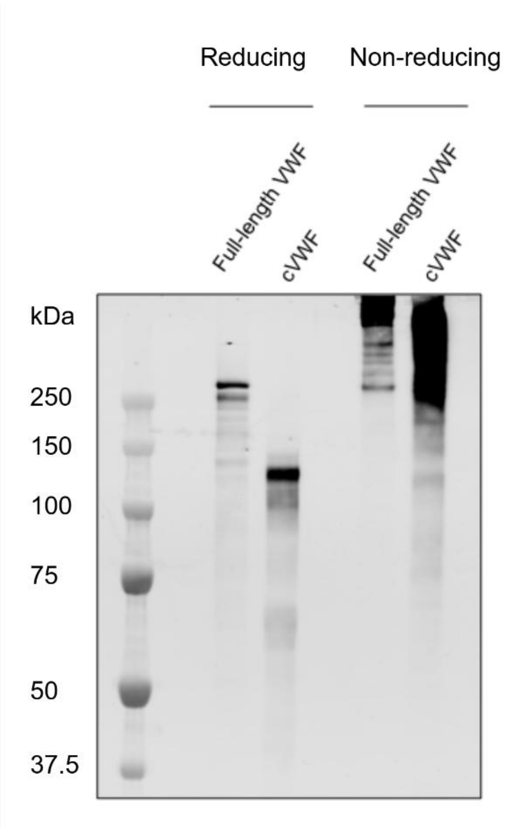


**Supplemental Figure 1. Western blot analysis of purified plasmin-cleaved VWF (cVWF) and intact VWF**. Purified cVWF was generated as previously described (El Otmani et al., Blood 2024). Under reducing conditions, intact VWF migrates at ~250 kDa, corresponding to the size of a VWF monomer. In cVWF, this band is no longer detected and distinct fragments appear at ~140, 120, and 70 kDa, confirming that each monomer has been cleaved at least once. Under non-reducing conditions, this shift in migration pattern is also evident: while intact VWF appears as a high molecular weight smear >250 kDa due to its polymeric nature, cVWF migrates as a broader smear spanning ~250 to 150 kDa, with an additional fragment at ~120 kDa. These findings indicate that cVWF retains a disulfide-linked polymeric structure despite proteolysis. Data are representative of at least three independent experiments.

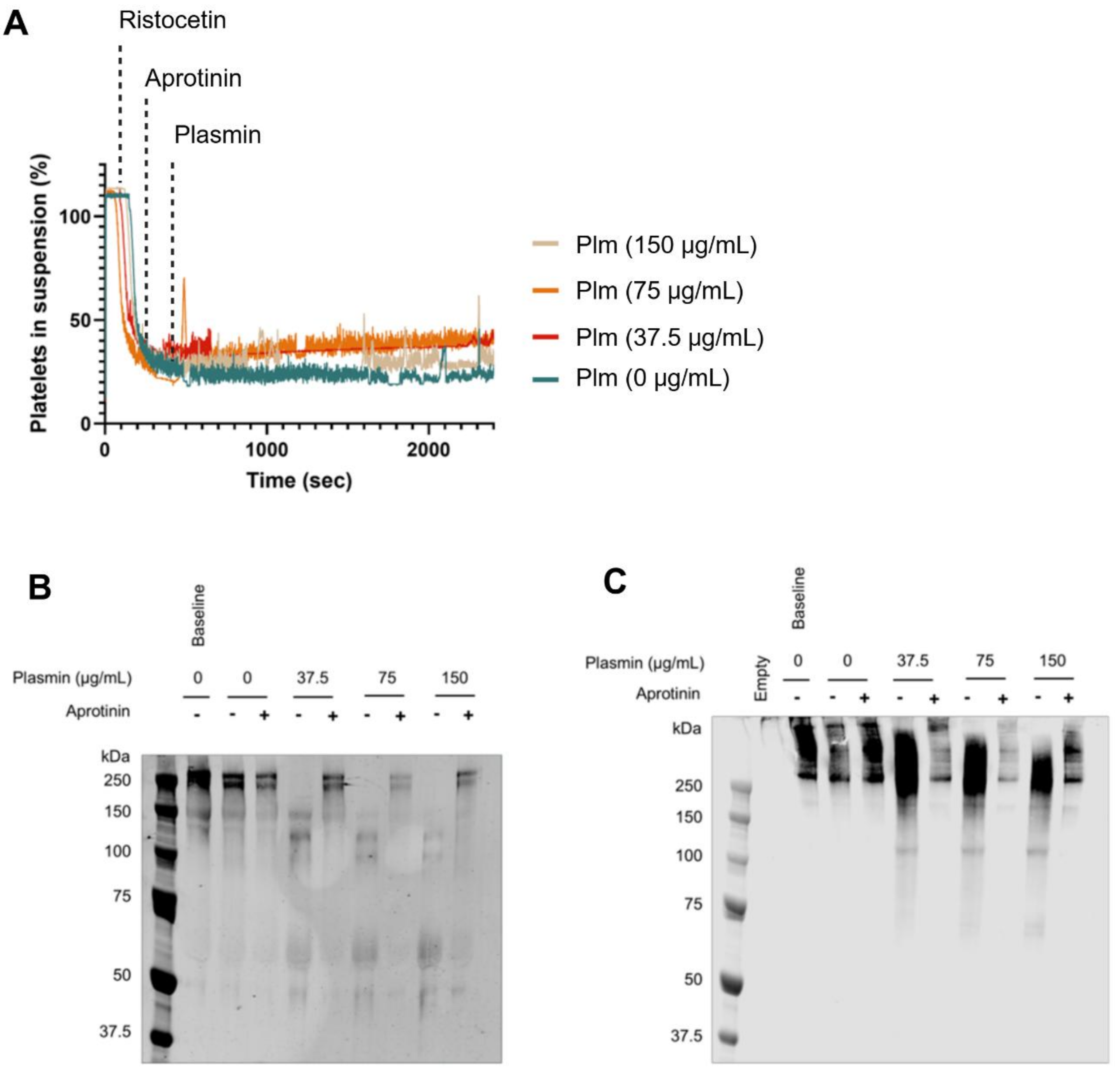


**Supplemental Figure 2. Plasmin cleaves VWF within agglutinates and is inhibited by aprotinin.** (A) Aprotinin (150 KIU/mL) effectively inhibited plasmin-mediated agglutinate breakdown. VWF in platelet-free supernatants from agglutinates treated with indicated plasmin concentrations for 30 minutes was analyzed by Western blot under (B) reducing and (C) non-reducing conditions. Increasing plasmin concentrations led to a progressive loss of high molecular weight VWF and the appearance of low molecular weight fragments, which was effectively blocked by aprotinin.

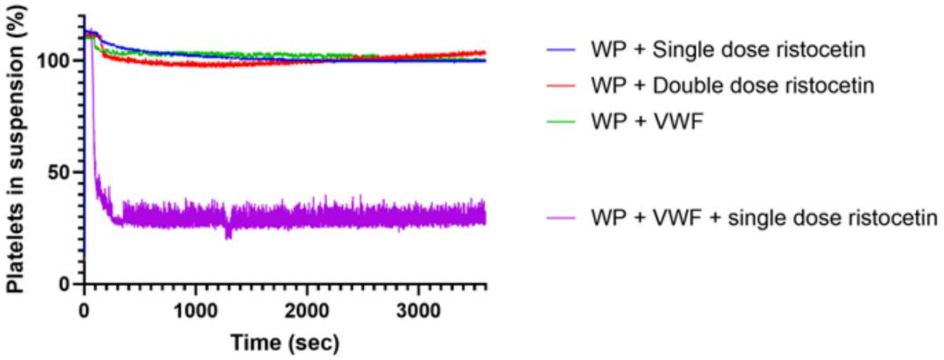


**Supplemental Figure 3. Double doses of ristocetin do not trigger spontaneous agglutinate formation.** In the absence of VWF, washed platelets subjected to a single (800 µg/mL) or a double dose of ristocetin (1600 µg/mL) do not form agglutinates. Adding VWF to washed platelets without a ristocetin trigger does not lead to agglutinate formation. A single dose of ristocetin induces agglutinates formation when washed platelets and VWF are combined. Data represent three independently executed experiments. WP = washed platelets.

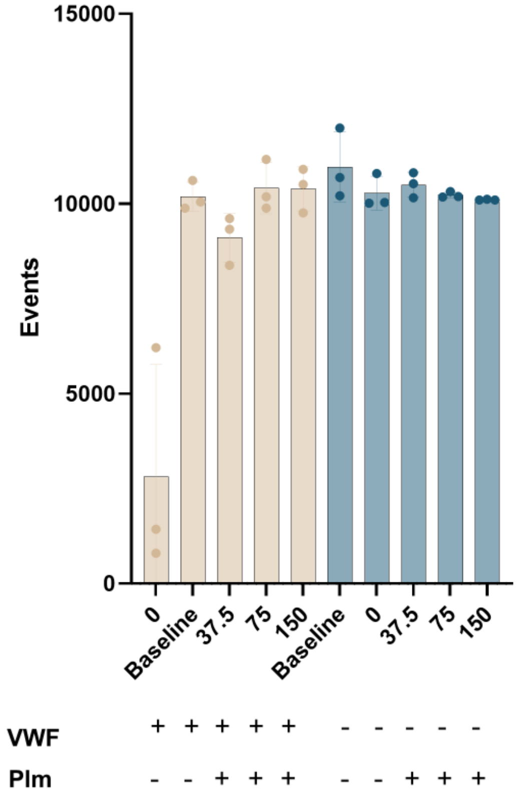


**Supplemental Figure 4. Event counts in flow cytometry experiments.** Insufficient single platelets could be recovered from experiments where agglutinates were not treated with plasmin, as most platelets remained in complex with VWF, thereby precluding reliable analysis.
